# FORGE-CRISPR: Reusable Modules for Focused CRISPR Library Construction

**DOI:** 10.64898/2026.06.15.732405

**Authors:** Ji-Ann Lee, Dylan Conklin, Michael Palazzolo, Jay M. Lee, Steven M. Dubinett

**Affiliations:** Division of Pulmonary, Critical Care and Sleep Medicine, Department of Medicine, David Geffen School of Medicine, University of California, Los Angeles, Los Angeles, CA, USA; Jonsson Comprehensive Cancer Center, University of California, Los Angeles, Los Angeles, CA, USA; Division of Thoracic Surgery, Department of Surgery, David Geffen School of Medicine, University of California, Los Angeles, Los Angeles, CA, USA

## Abstract

Pooled CRISPR screening has become a widely used approach for functional genomics, yet the construction of screening libraries remains tightly coupled to specific vector architectures and experimental formats. Incorporation of barcodes, multiplexed guide configurations, alternative fluorophores, selectable markers, or distinct screening modalities often requires reconstruction of entire libraries, even when the underlying biological content remains unchanged. These limitations are particularly acute in engineered cellular systems that already contain reporters, knock-in alleles, or pre-existing selection markers.

Here, we describe FORGE-CRISPR (Functional Oncology Research Genetic Engineering – CRISPR), a CRISPR library-construction system that separates biological guide content from screening-vector context. CRISPR knockout and CRISPR interference guide collections are first converted into reusable guide modules. Barcode modules and second-guide modules are generated separately and combined with guide modules during downstream assembly into screening acceptor vectors. This design allows guide-only, barcoded, and multiplexed libraries to be generated from shared physical components.

To support this framework, we developed a manufacturing workflow in which synthetic oligonucleotide pools are converted into reusable FORGE module libraries through PCR amplification, Golden Gate cloning, and background suppression. Using this approach, we constructed nine reusable guide-module libraries ranging from 45 to 4,830 guides (11,303 guides in total), and evaluated library quality through PCR–NGS analysis of guide representation and abundance distributions. As content for these libraries, we defined a modular, non-overlapping set of focused target collections, termed the druggable oncology genome, by partitioning druggable targets according to clinical development status and dependency distribution in DepMap, while drawing guide sequences from established, experimentally validated genome-wide libraries.

We report four resources: reusable guide, barcode, and second-guide modules; compatible screening acceptor vectors; a manufacturing and PCR–NGS quality-control workflow demonstrated across nine guide-module libraries; and a versioned set of focused druggable-oncology target collections.

## Introduction

Pooled CRISPR screening has become a central tool for functional genomics^1,2^, but the design requirements for different CRISPR modalities are not interchangeable. CRISPR knockout guides, CRISPRi guides, CRISPRa guides, and epigenome-editing guides are selected under different biological and positional constraints^3,4^. As a result, the value of a modular CRISPR library system is not that a single guide set can be used for every perturbation modality, but rather that different optimized guide collections can be manufactured, stored, and assembled using a common modular framework.

At the same time, screening applications increasingly require library formats that extend beyond simple guide-only vectors. Barcoded guide libraries can provide internal replicate structure, lineage-level information, and improved statistical power^5,6^. Schmierer and colleagues demonstrated that random sequence labels linked to guides can improve the accuracy and precision of CRISPR screen hit calling and enable robust analysis at reduced effective sampling depth^7^. This is especially important for focused libraries, primary cells, engineered reporter systems, and other settings in which cell number or experimental scale is limiting.

Despite the power of these approaches, incorporating barcodes, second guides, or alternative screening contexts into pooled CRISPR libraries remains technically cumbersome. In conventional workflows, guide selection, barcode incorporation, multiplex guide design, fluorophore choice, drug selection, and vector backbone are often coupled in a single monolithic library construction step. Consequently, changing one feature frequently requires rebuilding the full library, even when the biological design is otherwise unchanged.

This problem is particularly acute in engineered cell systems. Many experimental models already contain fluorescent reporters, endogenous knock-ins, selectable markers, or lineage-tracing elements. These pre-existing features constrain which CRISPR screening vectors can be used. A GFP-based vector may be incompatible with a GFP knock-in reporter; an mScarlet reporter may require a different fluorophore; an existing antibiotic cassette may restrict selectable-marker choice. Thus, practical screening often requires the same type of library architecture to be available in multiple vector contexts.

Here, we describe FORGE-CRISPR, a modular CRISPR library construction framework that separates guide selection, barcode incorporation, multiplexing, and screening-vector context into reusable physical components. Distinct CRISPRko and CRISPRi guide collections are converted into reusable FORGE guide modules. Barcode and second-guide modules can be generated independently and combined with guide modules as needed. These modules are then assembled into FORGE acceptor vectors that define the final screening context.

The system is designed to preserve the advantages of established CRISPR screening backbones while changing how libraries are manufactured and reused. Rather than introducing a new screening modality, this work introduces a modular construction strategy that allows focused guide collections, barcode architectures, and multiplexed guide formats to be built once, quality controlled, archived, and redeployed into multiple compatible screening vectors.

We further developed a workflow in which synthetic oligonucleotide pools are converted into reusable FORGE modules through PCR amplification, Golden Gate cloning, and background suppression, and applied this strategy to construct nine reusable guide-module libraries whose representation was validated at the module-library level by PCR–NGS. To define the biological content of these focused libraries, we partitioned druggable targets by clinical development status and by dependency distribution in DepMap^8–10^ to build a modular, non-overlapping target set, the druggable oncology genome, using guides drawn from established, experimentally validated genome-wide collections rather than designed de novo.

Together, FORGE-CRISPR applies modular cloning principles^11^ to pooled CRISPR library construction. It enables independent optimization of guide content, barcode architecture, multiplexing, and vector context while avoiding repeated reconstruction of complete libraries from the beginning.

The present study focuses on the design, implementation, manufacturing workflow, and quality control of the FORGE-CRISPR platform. Specifically, we describe the reusable module architecture, associated screening acceptor vectors, construction of reusable guide-module libraries from synthetic oligonucleotide pools, and PCR–NGS-based assessment of library representation and fidelity. The current work establishes the modular framework, reagent set, and library construction workflow. Cellular screening performance, guide-barcode benchmarking, and multiplexed-guide benchmarking are outside the scope of this preprint.

## Results

### Design rationale for a modular CRISPR library architecture

Focused CRISPR libraries have become increasingly important for studying specific biological processes, signaling pathways, and regulatory networks. However, conventional library construction tightly couples biological library content with the screening vector architecture. As a result, each new screening context, including CRISPR knockout, CRISPR interference, barcoded screening, multiplexed perturbation, or cell lines containing pre-existing reporters and selectable markers, often requires construction of an entirely new library from the beginning (Figure 1A).

**Figure 1.**
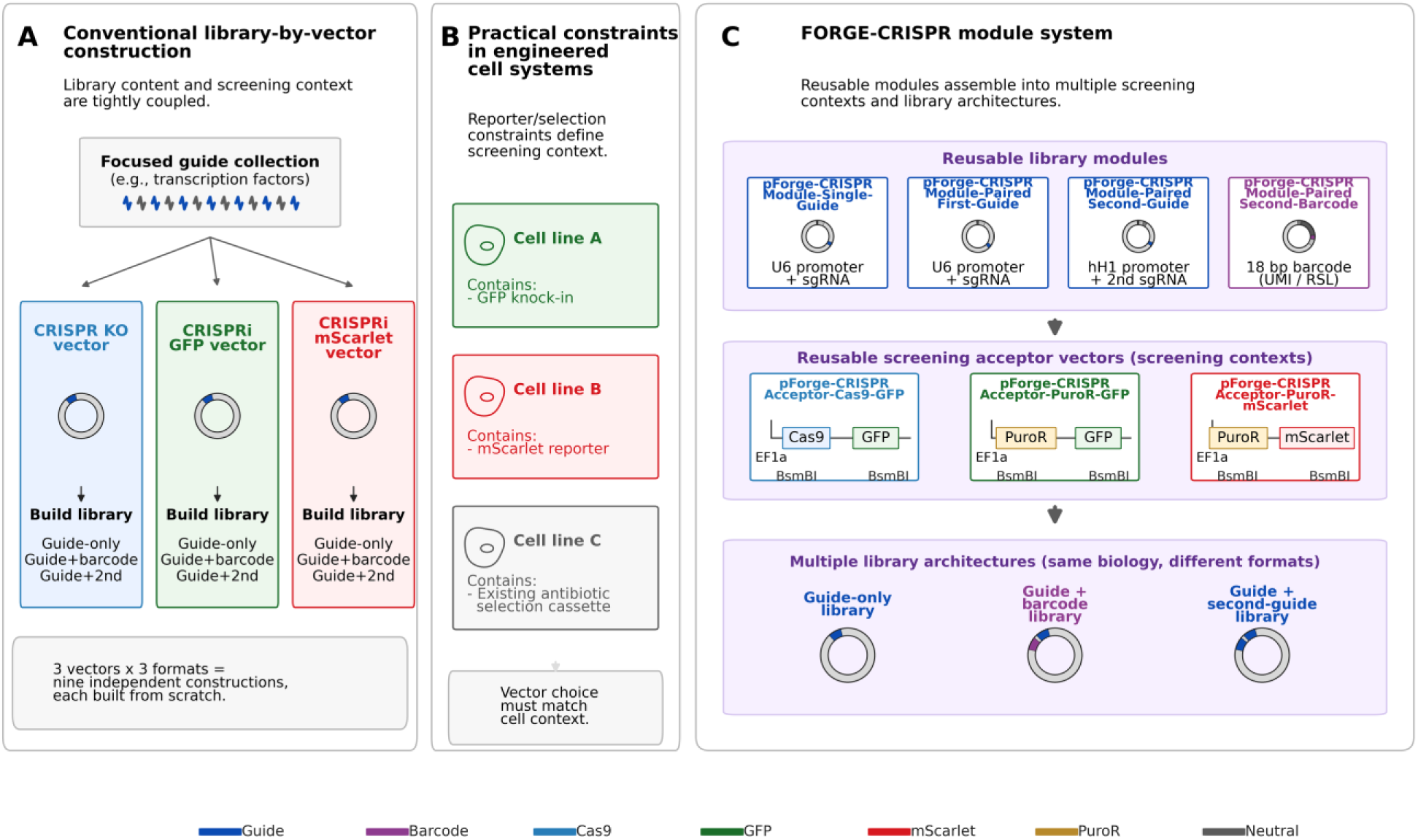
Design rationale for a modular CRISPR library architecture. (A) In the conventional, monolithic approach, biological library content and screening context are tightly coupled, so every combination requires independent library construction from the beginning. (B) Practical constraints in engineered cell systems: pre-existing reporters and selection cassettes force vector choice to be adapted to each cell line. (C) The FORGE-CRISPR modular solution: reusable guide, barcode, and second-guide modules assemble into multiple screening acceptor contexts and library architectures from a common set of components.

This limitation becomes particularly important in engineered cellular systems. Many experimental models already contain fluorescent reporters, drug-resistance markers, knock-in alleles, or other genetic modifications that constrain vector design. Consequently, the same biological guide collection may need to be rebuilt repeatedly in different vector formats to accommodate specific experimental requirements (Figure 1B).

To address this challenge, we developed the FORGE-CRISPR modular library architecture, which separates biological library content from screening context. In this framework, guide collections, barcode elements, and second-guide modules are generated as reusable components that can be assembled into multiple screening acceptor vectors. The resulting system enables rapid generation of guide-only, barcoded, and multiplexed library formats while preserving the underlying biological content (Figure 1C).

This design allows investigators to adapt library format to the experimental system without reconstructing the biological guide collection. As a result, the same guide content can be reused across multiple screening contexts, reducing cloning effort while increasing experimental flexibility.

### Architecture of reusable FORGE-CRISPR library modules and acceptor vectors

The FORGE-CRISPR system consists of four reusable library modules and three screening acceptor vectors (Figure 2). The reusable modules define the biological content of the library, whereas the acceptor vectors define the screening context. All constructs are deposited at Addgene under their pForge deposit names; internal identifiers (p-numbers) used in the figures are mapped to deposit names in Table 1.

**Figure 2.**
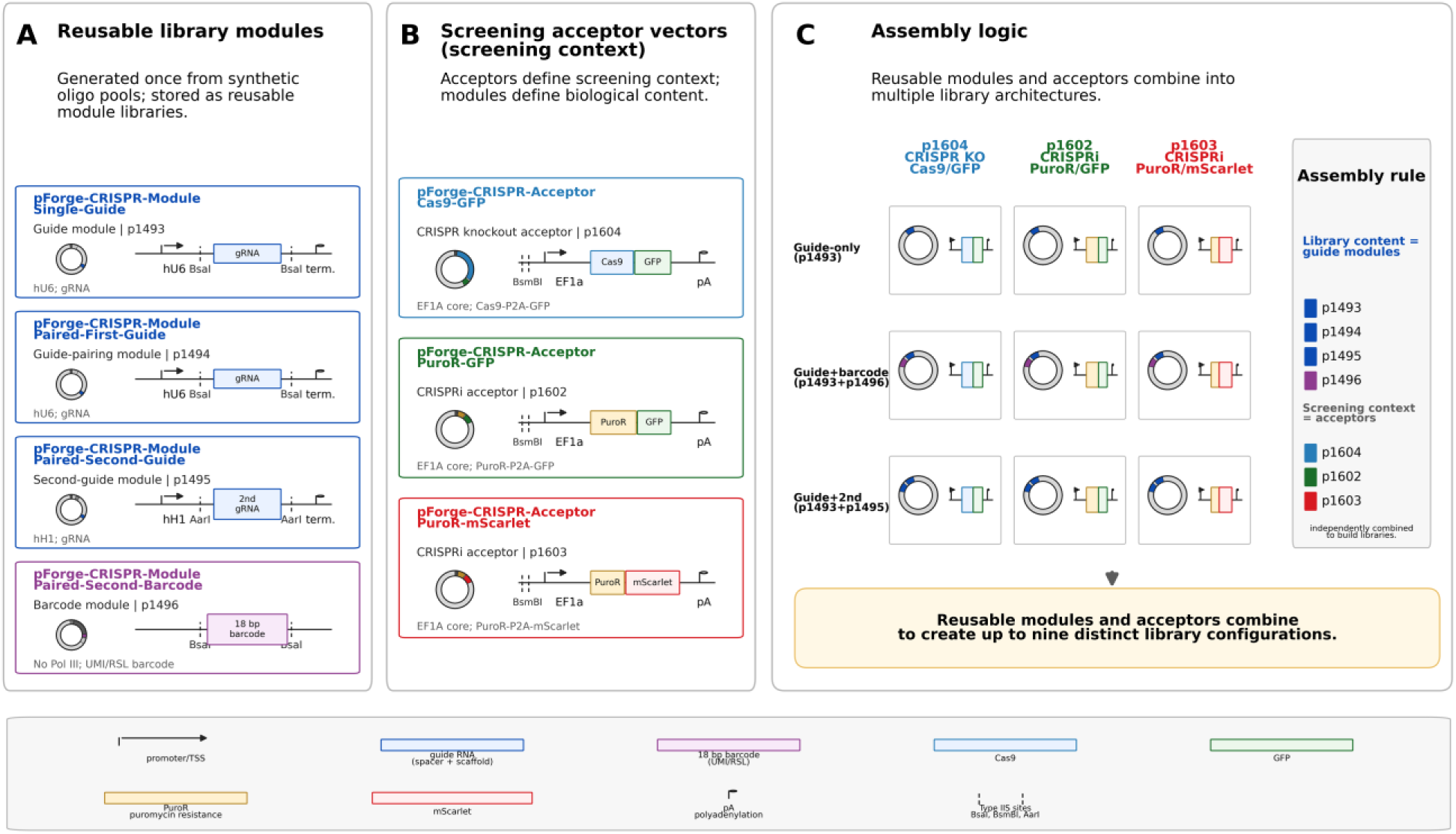
Architecture of reusable FORGE-CRISPR library modules and acceptor vectors. (A) Four reusable library modules (pForge deposit names with internal p-numbers), each maintained as a reusable plasmid library. (B) Three screening acceptor vectors defining knockout (Cas9/GFP) and CRISPRi (PuroR/GFP and PuroR/mScarlet) contexts; internal cloning uses BsaI/AarI and module insertion uses BsmBI. (C) Assembly logic: three library architectures combined with three acceptors yield up to nine distinct library configurations from a shared set of modules. Plasmid identifiers are listed in Table 1.

**Table 1.**
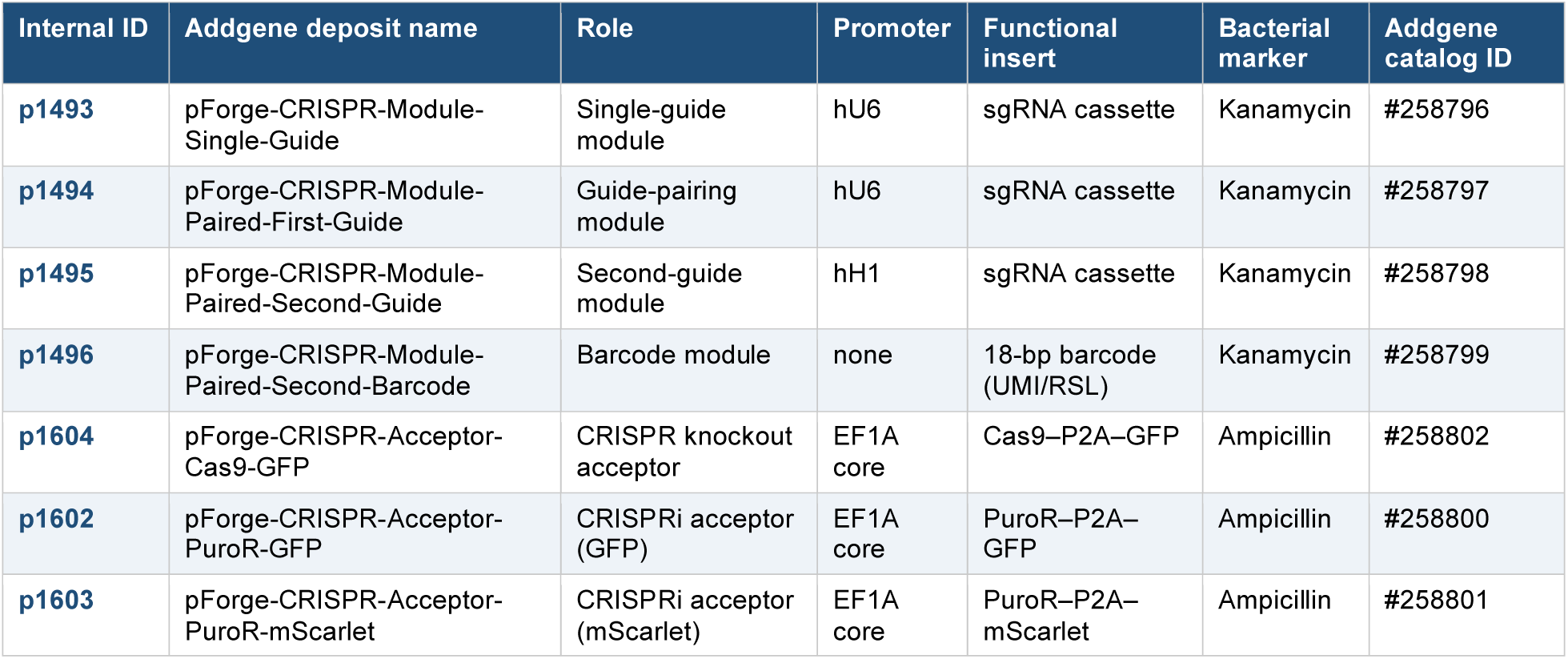
FORGE-CRISPR reusable modules and screening acceptor vectors. Addgene catalog IDs are listed for each construct (#258796–#258802).

| Internal ID | Addgene deposit name | Role | Promoter | Functional insert | Bacterial marker | Addgene catalog ID |
| --- | --- | --- | --- | --- | --- | --- |
| **p1493** | pForge-CRISPR-Module-Single-Guide | Single-guide module | hU6 | sgRNA cassette | Kanamycin | #258796 |
| **p1494** | pForge-CRISPR-Module-Paired-First-Guide | Guide-pairing module | hU6 | sgRNA cassette | Kanamycin | #258797 |
| **p1495** | pForge-CRISPR-Module-Paired-Second-Guide | Second-guide module | hH1 | sgRNA cassette | Kanamycin | #258798 |
| **p1496** | pForge-CRISPR-Module-Paired-Second-Barcode | Barcode module | none | 18-bp barcode (UMI/RSL) | Kanamycin | #258799 |
| **p1604** | pForge-CRISPR-Acceptor-Cas9-GFP | CRISPR knockout acceptor | EF1A core | Cas9-P2A-GFP | Ampicillin | #258802 |
| **p1602** | pForge-CRISPR-Acceptor-PuroR-GFP | CRISPRi acceptor (GFP) | EF1A core | PuroR-P2A-GFP | Ampicillin | #258800 |
| **p1603** | pForge-CRISPR-Acceptor-PuroR-mScarlet | CRISPRi acceptor (mScarlet) | EF1A core | PuroR-P2A-mScarlet | Ampicillin | #258801 |

The reusable modules include a single-guide module containing one sgRNA expression cassette (pForge-CRISPR-Module-Single-Guide; p1493), a guide-pairing module (pForge-CRISPR-Module-Paired-First-Guide; p1494), a second-guide module for multiplexed applications (pForge-CRISPR-Module-Paired-Second-Guide; p1495), and a barcode module containing an 18-bp identifier sequence (pForge-CRISPR-Module-Paired-Second-Barcode; p1496). Each module is generated independently and maintained as a reusable plasmid library (Figure 2A).

To support different screening applications, we generated three acceptor vectors. The knockout acceptor (pForge-CRISPR-Acceptor-Cas9-GFP; p1604) is an all-in-one CRISPR knockout vector containing Cas9 and GFP. The CRISPRi acceptor pForge-CRISPR-Acceptor-PuroR-GFP (p1602) supports CRISPR interference screening using a GFP-based selection scheme in cells expressing dCas9-KRAB^12,13^. The CRISPRi acceptor pForge-CRISPR-Acceptor-PuroR-mScarlet (p1603) provides an alternative CRISPRi format using mScarlet and puromycin selection (Figure 2B).

Assembly of reusable modules with screening acceptors generates multiple library architectures from a common set of biological components. Guide-only, guide-plus-barcode, and guide-plus-second-guide libraries can each be assembled into knockout or CRISPRi screening contexts, enabling up to nine distinct library configurations while preserving identical guide content (Figure 2C).

This separation allows the same guide content to be reused across acceptor vectors with different fluorophores, selection markers, and perturbation formats.

### Construction of reusable FORGE-CRISPR modules from synthetic oligonucleotide pools

To simplify library synthesis and reduce manufacturing costs, guide collections were encoded as synthetic oligonucleotide pools containing only guide-specific sequence information (Figure 3A). In this format, the oligonucleotide pool contains the biological content of the library but does not contain the additional sequence elements required for cloning, assembly, or screening applications^14^.

**Figure 3.**
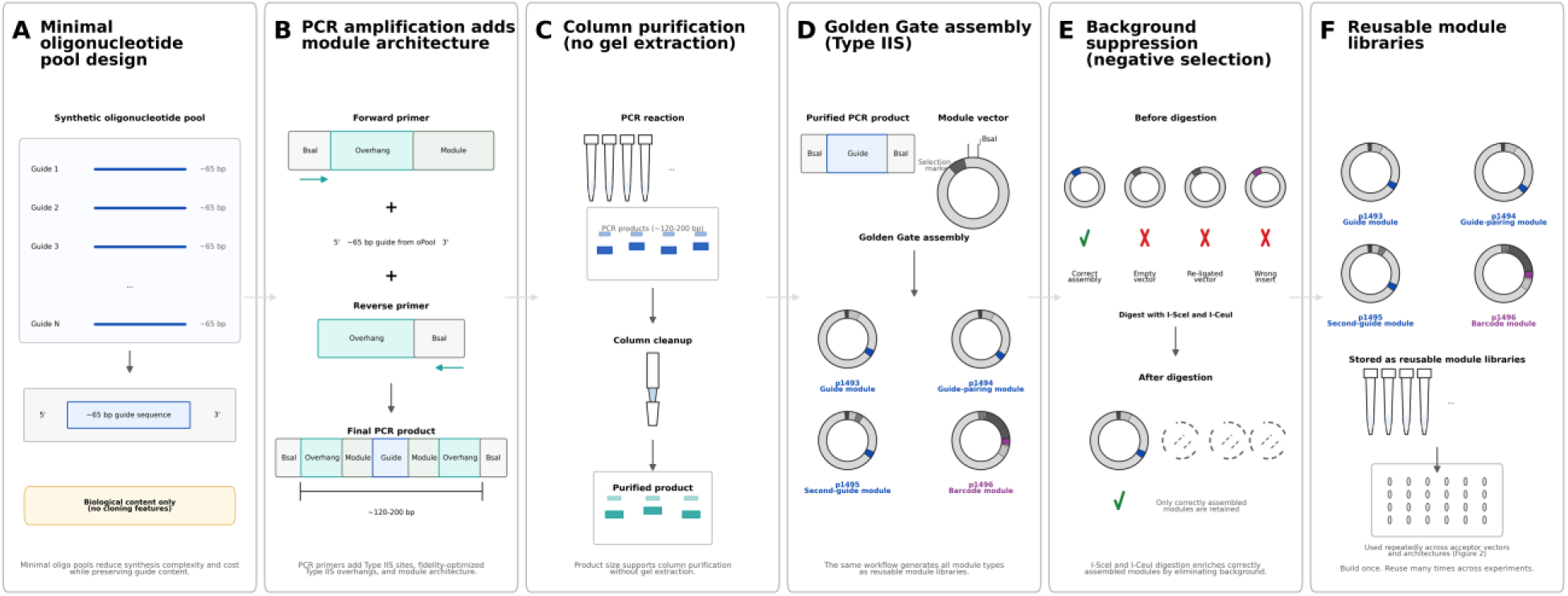
Construction of reusable FORGE-CRISPR modules from synthetic oligonucleotide pools. (A) Minimal oligonucleotide pool design encoding only guide content. (B) PCR amplification adds module architecture (Type IIS sites, fidelity-optimized Potapov overhangs^16^ (Potapov et al., 2018), and module-specific features). (C) Column purification without gel extraction. (D) Golden Gate assembly into module vectors. (E) Background suppression by I-SceI and I-CeuI negative selection. (F) Reusable module libraries archived for repeated assembly.

Module-specific architecture was introduced during PCR amplification through the use of primers containing Type IIS cloning sites, fidelity-optimized assembly overhangs, and module-specific sequence features (Figure 3B). This strategy converted minimal oligonucleotide pools into module-specific PCR products while maintaining a common manufacturing workflow across guide, barcode, and second-guide modules.

PCR amplification generated products of approximately 130 bp, enabling purification by standard column-based methods without gel extraction (Figure 3C). Purified PCR products were subsequently cloned into module vectors using Golden Gate assembly^15^ to generate reusable plasmid libraries corresponding to the desired module type (Figure 3D).

To reduce background arising from empty vectors, vector re-ligation, or incorrect assembly products, FORGE module vectors were designed to include negative-selection sites recognized by I-SceI and I-CeuI endonucleases; this digestion step is intended to eliminate undesired assembly products and enrich for correctly assembled modules (Figure 3E).

Using this workflow, guide modules, guide-pairing modules, second-guide modules, and barcode modules were generated through a common manufacturing process and maintained as reusable plasmid libraries (Figure 3F). Once constructed, these modules could be repeatedly assembled into multiple screening architectures without reconstruction of the underlying biological library content.

### Construction and PCR–NGS validation of independent modular CRISPR libraries

To evaluate the scalability of the FORGE-CRISPR cloning framework, nine focused CRISPR guide libraries were constructed from pooled synthetic oligonucleotides. Modules 1–5 contained 138, 528, 738, 1,650, and 204 guides, respectively. Additional focused libraries included Module 6 (45 guides) and Module 7 (152 guides), while larger libraries were generated in Module 10 (3,018 guides) and Module 11 (4,830 guides). Collectively, these libraries comprised 11,303 unique guide RNAs spanning a broad range of library complexities.

The guide libraries were synthesized as pooled oligonucleotides containing a guide sequence flanked by constant priming handles. For Modules 1–7, pooled oligonucleotides were resuspended to generate master stocks corresponding to approximately 10 nM per guide, with resuspension volumes scaled according to library size and oligonucleotide yield. Working stocks were generated by dilution to approximately 1 nM per guide and used directly as templates for library amplification.

Library amplification was performed using multiple parallel PCR reactions to minimize amplification bias and preserve representation of low-abundance library members, with the number of parallel reactions adjusted according to library complexity^17^. The PCR primers introduced BsaI recognition sites and assembly overhangs required for modular Golden Gate cloning; amplification of the approximately 60 bp guide templates generated products of approximately 130 bp, allowing efficient separation of amplified products from residual primers by silica spin-column purification without gel extraction.

Purified guide libraries were cloned by BsaI Golden Gate assembly into guide module vectors. Two guide module formats were developed: the single-guide module (p1493) generated single-guide libraries that could be transferred directly into downstream screening vectors, whereas the guide-pairing module (p1494) generated guide libraries that could subsequently be combined with either barcode modules (p1496) or second-guide modules (p1495) to generate guide-barcode or multiplexed guide configurations^18,19^. In all cases, guide libraries were first converted into reusable module libraries prior to assembly into screening vectors.

Following transformation, library complexity was assessed by titration plating and used to determine expansion scale. Across all libraries, transformant numbers substantially exceeded library complexity, providing hundreds to thousands of transformants per guide and minimizing representation loss during amplification. Guide modules will be subsequently assembled into a family of lentiviral FORGE acceptor vectors using BsmBI-mediated Golden Gate assembly, supporting CRISPR knockout, CRISPRi, guide-barcode, and multiplexed guide configurations while preserving a common guide library architecture^5^.

To avoid ambiguity, we use three terms consistently. A guide is the CRISPR targeting spacer that defines the biological content of a library. The barcode module contributes a fixed 18-bp identifier sequence, also referred to as a random sequence label. This barcode is encoded in the plasmid library and physically linked to a guide during assembly. By contrast, the sequencing UMI is a random 8-nucleotide sequence introduced during the first PCR of next-generation sequencing library preparation to label individual template molecules and support duplicate control. Thus, the “valid guide + UMI” category refers to reads in which both the guide spacer and a well-formed sequencing UMI were recovered.

Following module construction, library composition was evaluated by PCR amplification and next-generation sequencing (Figure 4). Across the nine module sublibraries, 99.3–100.0% of designed guides were detected, 96.7–98.6% of reads were valid (guide + UMI), mapping rates were 97.4–98.3%, and abundance skew was low (Gini 0.137–0.264), supporting broad guide recovery and relatively uniform representation across more than two orders of magnitude in library complexity.

**Figure 4.**
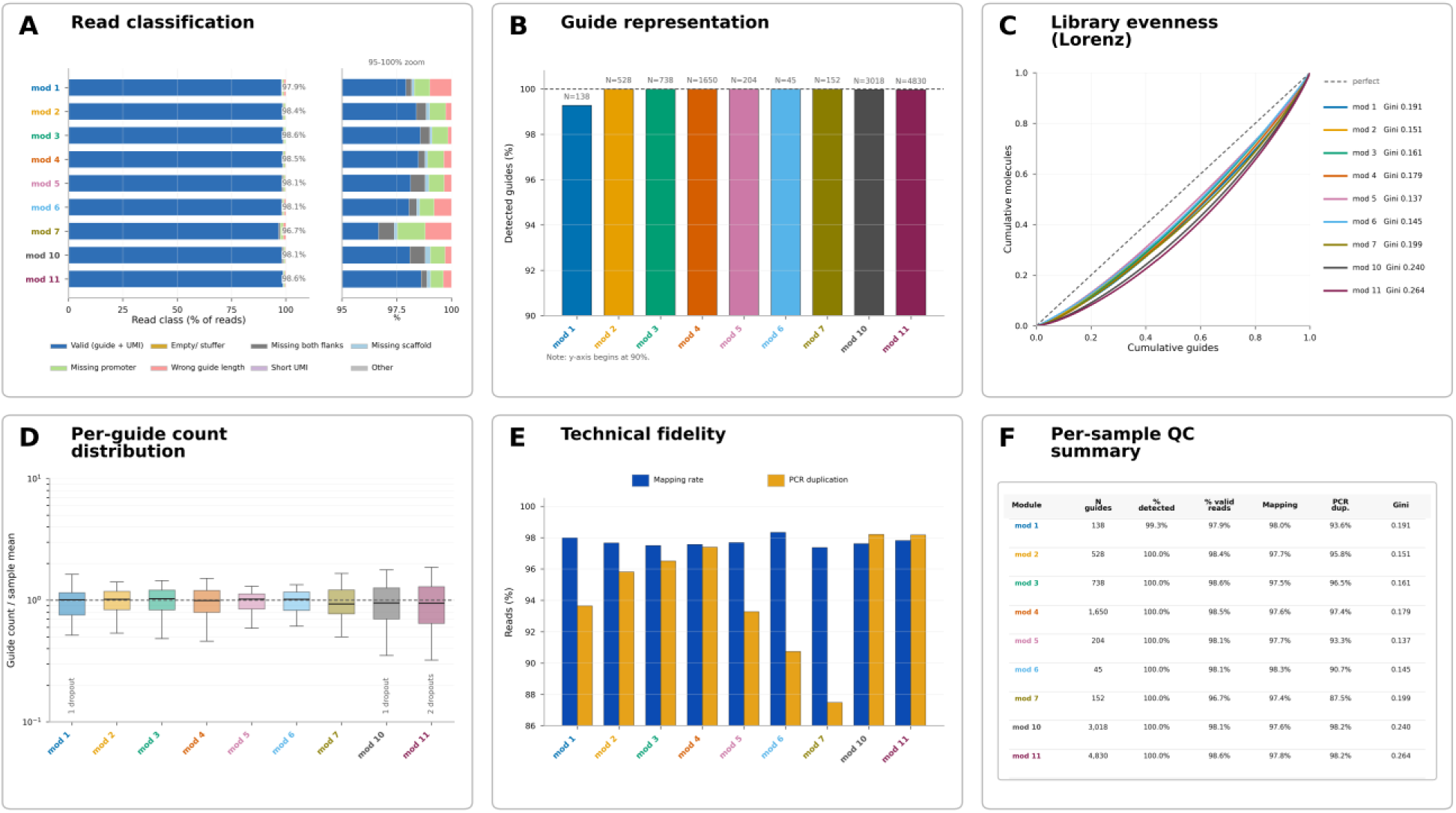
Sequencing quality control of the reusable FORGE-CRISPR module libraries. (A) Read classification per sublibrary (fraction of valid guide + UMI reads versus invalid categories). (B) Guide representation (percent of designed guides detected; library size annotated). (C) Library evenness shown as Lorenz curves with per-sample Gini coefficients. (D) Per-guide count distribution normalized to the sample mean. (E) Technical fidelity (mapping rate and PCR duplication). (F) Per-sample QC summary across the nine module sublibraries.

Collectively, these experiments establish a framework for systematic quality control of modular FORGE-CRISPR libraries and demonstrate that libraries generated through the modular workflow can be evaluated using standard PCR–NGS approaches prior to assembly into final screening vectors.

### Modular library assembly enables efficient generation of reusable CRISPR libraries

We designed the module system to enable repeated reuse of guide content without requiring reconstruction of complete screening libraries. The workflow was designed such that only a small fraction of the material generated at each stage was required to seed the subsequent step: synthetic oligonucleotide pools were amplified and cloned into reusable module vectors, and module libraries were then assembled into screening acceptor vectors to generate final CRISPR screening libraries. Because each stage produced substantially more material than was required for the next step, module libraries could be archived and reused for future library assembly (Figure 5).

**Figure 5.**
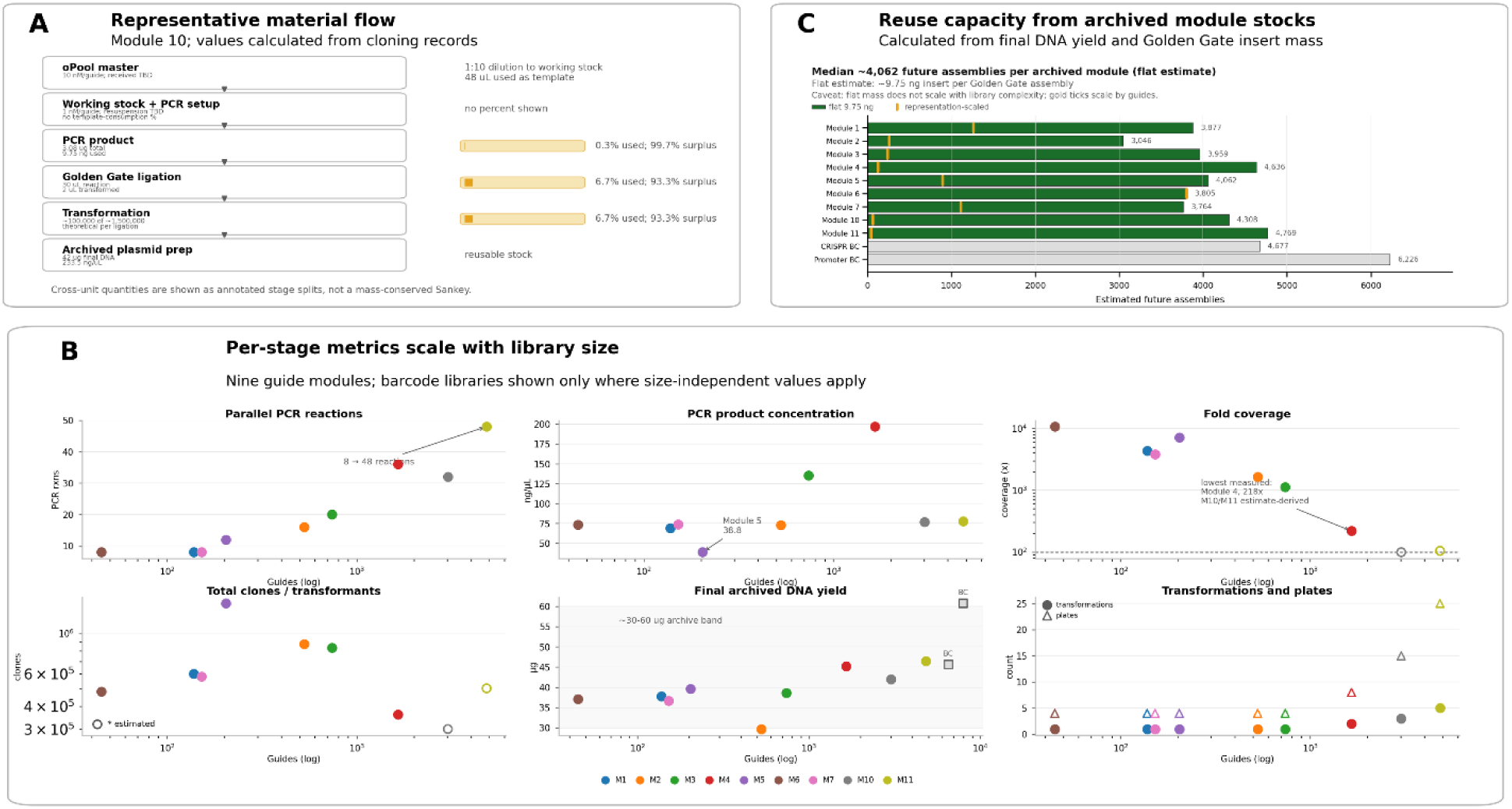
Material economy and reuse of the FORGE-CRISPR modular workflow. (A) Representative material flow for a single library (Module 10), with values parsed directly from the cloning records. At each stage only a small fraction of the available intermediate is consumed to seed the next step, leaving the remainder as archivable surplus: ∼9.75 ng of the ∼3.08 µg of purified PCR product (∼0.3%) is used per Golden Gate assembly, 2 µL of the 30 µL ligation (∼6.7%) is carried into transformation, and ∼100,000 of an estimated ∼1,500,000 possible transformants per ligation are recovered. Because the quantities span different units, each step is shown as an annotated used-versus-surplus split rather than a mass-conserved Sankey. (B) Per-stage metrics across the nine constructed guide modules as a function of library size (guides, log scale), with the two barcode libraries shown only for size-independent measures: number of parallel PCR reactions, PCR product concentration, fold representation (clone coverage; dashed reference at 100×, with Modules 10 and 11 shown as open symbols because their coverage is derived from estimated clone counts), total clones/transformants (open symbols denote estimated counts), final archived module-library DNA yield (∼30–60 µg across all libraries), and the number of transformations and expansion plates. (C) Reuse capacity of each archived module library, estimated as its final DNA yield divided by the ∼9.75 ng of insert consumed per Golden Gate assembly (median ≈ 4,062 future assemblies per module); the flat per-assembly estimate (green) is shown alongside a representation-scaled estimate (orange) that accounts for the greater insert mass required to preserve representation in higher-complexity libraries. Together these panels quantify the ‘build once, reuse many times’ principle: a small fraction of each cloning intermediate seeds the next step, while high-yield archived module stocks support repeated assembly into multiple screening contexts.

To quantify workflow performance, material use and library complexity were tracked across construction and related to library size, including PCR product yield and concentration, the number of parallel PCR reactions, clone coverage, and archived module-library DNA yield (Figure 5A,B). Dividing the archived DNA yield of each module by the small insert mass consumed per assembly indicates that a single archived module supports on the order of several thousand future library assemblies (Figure 5C). These measurements quantify workflow scalability across library sizes and show that guide-module construction can be separated from downstream acceptor-vector deployment.

### Definition of focused, non-overlapping libraries spanning the druggable oncology genome

We next defined focused libraries with compact, interpretable, and reusable gene content. Rather than designing guides de novo, we drew guide sequences from established, experimentally validated genome-wide collections and concentrated the design effort on defining which genes each focused library should contain (Figure 6). CRISPRi modules used the top six guides per transcript from the Weissman/Replogle hCRISPRi-v2.1 library, and CRISPR knockout modules used the top two guides per gene from the Gonçalves minimal CRISPR-Cas9 library (minilibcas9)^20^. Building modules from previously benchmarked guides means that these guides carry prior large-scale on-target and specificity characterization rather than untested predictions, although their performance in the FORGE library context remains to be evaluated empirically.

**Figure 6.**
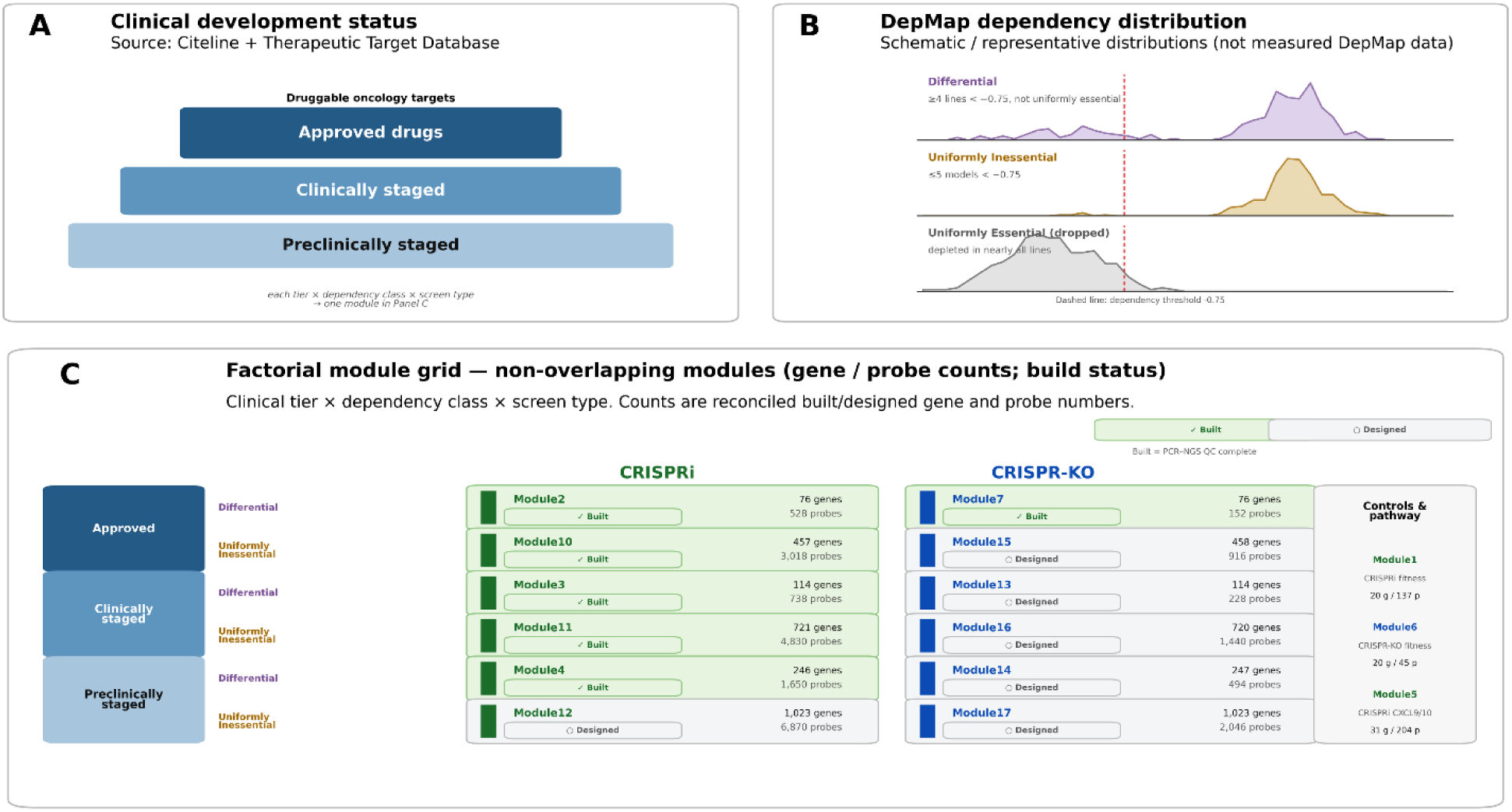
Definition of focused, non-overlapping libraries spanning the druggable oncology genome. (A) Druggable oncology targets stratified by clinical development status—targets of approved, clinically staged, or preclinically staged drugs—using Citeline and the Therapeutic Target Database. (B) Classification of each target’s dependency distribution across cancer cell lines (DepMap gene-effect score) into ‘differential’ (selectively required: ≥4 cell lines below −0.75 and not uniformly essential) and ‘uniformly inessential’ (≤5 models below −0.75); uniformly essential genes (depleted in nearly all lines) are excluded. The distributions shown are representative illustrations of each class rather than the underlying DepMap data. (C) The resulting factorial module grid (clinical tier × dependency class × screen type) of non-overlapping CRISPRi and CRISPR knockout modules, annotated with gene and probe counts and build status (validated by PCR–NGS versus designed-and-queued); the right-hand column lists the shared fitness-control modules and the focused CXCL9/CXCL10 pathway module that accompany every screen. CRISPRi modules draw the top six guides per transcript from the Weissman/Replogle hCRISPRi-v2.1 library and CRISPR knockout modules the top two guides per gene from the Gonçalves minilibcas9 library (Materials and Methods). Module identifiers are shared with the constructed sublibraries in Figures 4 and 5 and with Table 2.

**Table 2.** Build status and PCR–NGS quality control of the FORGE-CRISPR module libraries. The nine constructed guide-module libraries (11,303 guides in total) were each assessed by PCR–NGS for guide representation (percent of designed guides detected), the fraction of valid guide + UMI reads, mapping rate, PCR duplication, and abundance evenness (Gini coefficient). All nine were built as reusable plasmid module libraries and quality-controlled; the remaining cells of the clinical-tier × dependency-class × screen-type grid are defined and queued for construction but have not yet been built or sequenced. Module identifiers are shared with Figures 4–6.

| Module | Guides (designed) | Detected (%) | Valid reads (%) | Mapping (%) | Duplication (%) | Gini | Build status |
| --- | --- | --- | --- | --- | --- | --- | --- |
| 1 | 138 | 99.3 | 97.9 | 98.0 | 93.6 | 0.191 | Built; PCR-NGS QC complete |
| 2 | 528 | 100.0 | 98.4 | 97.7 | 95.8 | 0.151 | Built; PCR-NGS QC complete |
| 3 | 738 | 100.0 | 98.6 | 97.5 | 96.5 | 0.161 | Built; PCR-NGS QC complete |
| 4 | 1,650 | 100.0 | 98.5 | 97.6 | 97.4 | 0.179 | Built; PCR-NGS QC complete |
| 5 | 204 | 100.0 | 98.1 | 97.7 | 93.3 | 0.137 | Built; PCR-NGS QC complete |
| 6 | 45 | 100.0 | 98.1 | 98.3 | 90.7 | 0.145 | Built; PCR-NGS QC complete |
| 7 | 152 | 100.0 | 96.7 | 97.4 | 87.5 | 0.199 | Built; PCR-NGS QC complete |
| 10 | 3,018 | 100.0 | 98.1 | 97.6 | 98.2 | 0.240 | Built; PCR-NGS QC complete |
| 11 | 4,830 | 100.0 | 98.6 | 97.8 | 98.2 | 0.264 | Built; PCR-NGS QC complete |

The main content-design step is a modular, non-overlapping partition of the druggable genome, which we refer to as the druggable oncology genome. Target druggability and clinical development status were obtained from commercial and public drug-target resources (Citeline and the Therapeutic Target Database^21^), and each target was assigned to one of three clinical-status tiers: targets of approved drugs, of clinically staged drugs, or of preclinically staged drugs (Figure 6A). In parallel, each target’s dependency distribution across cancer cell lines was classified using DepMap. Targets were classified as differential when at least four cell lines had a dependency score below −0.75 and the target was not uniformly essential. Targets were classified as uniformly inessential when five or fewer models crossed this threshold (Figure 6B). Combining the three clinical tiers with the two dependency classes, separately for CRISPRi and CRISPR knockout, defines a factorial grid of non-overlapping modules in which each target is assigned to exactly one module per screen type.

Applying this scheme produced a set of focused modules spanning more than two orders of magnitude in size, from tens to roughly one thousand genes per module (Figure 6C; Table 2). A representative subset has been constructed and validated by PCR–NGS, including the approved-, clinical-, and preclinical-tier differential CRISPRi modules and the approved- and clinical-tier uniformly inessential CRISPRi modules, together with the approved-tier differential knockout module; the remaining grid cells are defined and queued for construction. Because the gene sets are defined independently of screen type, the CRISPRi and CRISPR knockout versions of each module target the same genes: across the defined modules the two formats agreed to within a single gene per cell, with the rare differences attributable to individual targets lacking a usable guide in one of the two source libraries (for example, GRK2 and CHRM3 lack a CRISPRi guide in the source library, and UGT1A1 lacks a knockout guide).

Two additional module classes complement the druggable oncology genome. Fitness-control modules contain ten essential and ten inessential genes^22,23^ selected for consistent behavior across multiple CRISPR knockout and CRISPRi screens and distributed across the genome, providing internal performance standards for every screen (Figure 6C, right). A focused biologic-pathway module targeting genes implicated in CXCL9/CXCL10 regulation (Module 5) illustrates how pathway-specific collections are assembled within the same framework (Figure 6C, right).

This strategy converts large public druggability and dependency resources into compact, non-overlapping CRISPR modules built from validated guides. By partitioning targets along clinically and biologically meaningful axes, the resulting modules can be screened individually or combined. Because the modules share the FORGE-CRISPR architecture, they can be reformatted across knockout and CRISPRi contexts without redesign.

## Discussion

The central objective of this work was not to develop a new CRISPR screening modality, but rather to introduce FORGE-CRISPR, a modular framework for CRISPR library construction. Current CRISPR library workflows frequently couple guide selection, barcode architecture, multiplexing strategy, fluorophore selection, drug resistance markers, and vector backbone into a single library construction process. As a consequence, changes in screening format often require reconstruction of the entire library, even when the underlying biological content remains unchanged. Accordingly, this preprint establishes the platform’s design, reusable reagent set, manufacturing workflow, and module-level PCR–NGS quality control, rather than benchmarking screening performance in cells.

The modular architecture described here separates these functions into reusable components. Guide collections, barcode modules, multiplexing modules, and screening acceptors can be independently designed, manufactured, validated, and archived. This separation enables focused guide collections to be deployed in multiple experimental contexts without repeatedly rebuilding complete libraries from the beginning. The system preserves compatibility with established CRISPR screening architectures while introducing flexibility in how libraries are constructed and reused.

The increasing availability of large public CRISPR screening datasets and dependency maps^8,10^ has also changed how screening libraries are designed. Rather than constructing increasingly large libraries, many biological questions can now be addressed using focused collections targeting specific pathways, transcriptional regulators, kinases, epigenetic factors, or context-dependent dependencies. We therefore defined focused content by partitioning druggable targets along their dependency distribution in DepMap and their clinical development status, drawing guides from established, experimentally validated genome-wide libraries, so that compact, information-rich modules could be assembled without redesigning guides. In this setting, modular library construction becomes particularly valuable because focused libraries are frequently redesigned, expanded, and adapted as new biological information becomes available.

An additional motivation for this work was the growing importance of barcode-based screening strategies. Molecular barcodes can provide internal replicate structure, improve statistical power, and enable more sophisticated analysis of pooled screening experiments. However, barcode incorporation often requires construction of specialized libraries that are difficult to modify or repurpose. By separating barcode content from guide content, the FORGE-CRISPR architecture allows barcode-containing and non-barcoded libraries to be generated from related module sets while preserving flexibility in downstream assembly.

This study has several limitations. First, validation is performed at the level of plasmid-library construction and PCR–NGS representation rather than pooled cellular screen performance; the libraries have not yet been transduced, selected, or screened in cells. Second, although the architecture is designed to support guide-only, barcoded, and second-guide configurations across multiple acceptor contexts, the present data do not benchmark each final screening-vector format, and only a representative subset of acceptor assemblies has been generated. Third, the focused druggable-oncology modules depend on specific versions of DepMap, Citeline, the Therapeutic Target Database, and the source guide libraries, and future releases may alter target membership; the partition is therefore best treated as a versioned, reproducible design resource rather than a fixed biological taxonomy. Fourth, the current implementation covers CRISPR knockout and CRISPR interference, and additional modalities will require modality-specific guide design and acceptor validation.

The current implementation has focused on CRISPR knockout and CRISPR interference screening. However, the modular principles underlying FORGE-CRISPR are not specific to these applications. Future implementations could incorporate guide collections optimized for CRISPR activation, CRISPRoff, alternative Cas nucleases, base editors, prime editors, or other emerging perturbation technologies^25–28^. Similarly, additional acceptor vectors could be developed to support specialized reporters, selectable markers, or cell-type-specific screening requirements. These extensions would not require redesign of the overall framework, but rather the addition of new module classes and compatible acceptor architectures.

## Materials and Methods

### Construction strategy for FORGE-CRISPR modular libraries

FORGE-CRISPR libraries were constructed using a modular cloning workflow in which pooled guide, barcode, and second-guide elements were first converted into reusable plasmid module libraries and subsequently assembled into lentiviral screening acceptor vectors^29^. Internal plasmid identifiers are provided in parentheses. Reusable module vectors included pForge-CRISPR-Module-Single-Guide (p1493), pForge-CRISPR-Module-Paired-First-Guide (p1494), pForge-CRISPR-Module-Paired-Second-Guide (p1495), and pForge-CRISPR-Module-Paired-Second-Barcode (p1496). Screening acceptor vectors included pForge-CRISPR-Acceptor-PuroR-GFP (p1602), pForge-CRISPR-Acceptor-PuroR-mScarlet (p1603), and pForge-CRISPR-Acceptor-Cas9-GFP (p1604). The single-guide module was used to generate guide-only libraries, whereas the paired-guide module was used when guide libraries were subsequently combined with either barcode modules or second-guide modules. This design allowed a common guide library to be archived as a reusable module and reformatted into multiple screening contexts without reconstructing the guide pool.

### Synthetic oligonucleotide pool design and preparation

Guide libraries were synthesized as pooled oligonucleotides containing guide-specific sequences flanked by constant priming handles. Minimal oligonucleotide designs were used so that the synthetic pools encoded the biological library content, whereas cloning sites, assembly overhangs, and module-specific sequence features were introduced during PCR amplification. Guide pools for Modules 1, 6, and 7 were obtained as IDT oPools (Integrated DNA Technologies), and guide pools for Modules 2, 3, 4, 5, 10, and 11 were obtained as Twist Bioscience oligo pools (Twist Bioscience). Modules 1–5 contained 138, 528, 738, 1,650, and 204 guides, respectively; Modules 6 and 7 contained 45 and 152 guides, respectively; and Modules 10 and 11 contained 3,018 and 4,830 guides, respectively.

For guide libraries, oligonucleotide pools were resuspended to generate master stocks corresponding to approximately 10 nM per guide. For IDT pools specified as pmol per oligo, 1 pmol per guide was resuspended in 100 µL and 10 pmol per guide was resuspended in 1,000 µL. For Twist pools reported by mass, the amount per oligo was converted to pmol based on oligonucleotide length and then resuspended using the same rule. Working stocks were prepared by 1:10 dilution of master stocks to approximately 1 nM per guide and used directly as PCR templates.

### PCR amplification of pooled guide and barcode inserts

Pooled oligonucleotide libraries were amplified by multiple parallel PCR reactions to reduce amplification bias and minimize PCR jackpotting of individual guides. Each 10 µL PCR reaction contained 2.0 µL 5× Phusion HF buffer, 0.2 µL 10 mM dNTPs, 0.5 µL 10 µM forward primer, 0.5 µL 10 µM reverse primer, 0.1 µL Phusion DNA polymerase (Thermo Scientific, F530), 6.2 µL nuclease-free water, and 1.5 µL of 1 nM-per-guide working stock. PCR cycling was performed with an initial denaturation at 98°C for 30 s; 12 cycles of 98°C for 10 s, 62°C for 15 s, and 72°C for 10 s; followed by a final extension at 72°C for 60 s and hold at 4°C.

The number of parallel PCR reactions was scaled according to library complexity. Libraries containing fewer than 200 guides were amplified using 8–12 parallel reactions; libraries containing 200–600 guides were amplified using 12–16 reactions; libraries containing 600–1,000 guides were amplified using 16–20 reactions; libraries containing 1,000–2,000 guides were amplified using 32–36 reactions; libraries of approximately 3,000 guides were amplified using approximately 32 reactions; and libraries of approximately 5,000 guides were amplified using approximately 48 reactions. After amplification, parallel reactions were pooled, a small aliquot was analyzed by agarose gel electrophoresis, and the remaining pooled PCR product was purified by silica spin-column purification and quantified.

Guide pools amplified with IDT0787 and IDT0788 generated products of approximately 131–132 bp, depending on the length of the guide-pool insert. Barcode oligonucleotides were amplified with IDT0789 and IDT0790 to generate a 156 bp product. The short amplicon sizes allowed removal of residual primers and small oligonucleotides by column purification without gel extraction.

### Golden Gate cloning of reusable guide modules

Purified guide PCR products were cloned into reusable guide module vectors by BsaI-mediated Golden Gate assembly (NEB, E1601S). Cleaned PCR inserts were diluted to approximately 7.5 ng/µL. For each 20 µL Golden Gate reaction, 75 ng of vector was combined with approximately 9.75 ng of insert, 2.0 µL 10× ligase buffer, 1.0 µL enzyme mix or restriction enzyme mix, and nuclease-free water to 20 µL. For an approximately 130 bp insert and an approximately 3.6 kb vector, this corresponded to an insert-to-vector molar ratio of approximately 3.6:1.

Single-guide libraries were cloned into pForge-CRISPR-Module-Single-Guide (p1493) when the guide library was intended to be transferred directly into downstream acceptor vectors without a barcode or second guide. Paired-guide libraries were cloned into pForge-CRISPR-Module-Paired-First-Guide (p1494) when the guide library was intended for later combination with either a barcode module or a second-guide module. The resulting guide-module plasmid pools were maintained as reusable plasmid libraries.

### Golden Gate cloning of reusable barcode modules

Barcode modules were generated by PCR amplification of barcode oligonucleotides followed by BsaI-mediated Golden Gate cloning into pForge-CRISPR-Module-Paired-Second-Barcode (p1496). The barcode module was designed to provide an 18-bp identifier sequence that could be paired with a guide module during downstream assembly into an acceptor vector. Where indicated, a zero-barcode control oligonucleotide was used as a defined barcode-control insert. Barcode module libraries were transformed, expanded, and archived in the same manner as guide-module libraries.

### Construction of second-guide modules

Second-guide modules were generated in pForge-CRISPR-Module-Paired-Second-Guide (p1495). For defined second-guide constructs, complementary 5′-phosphorylated oligonucleotides were annealed to generate double-stranded guide inserts with compatible overhangs and cloned into p1495 using AarI-mediated Type IIS assembly. For the KRAS second-guide construct, 5′-phosphorylated oligonucleotides IDT0813 and IDT0814 were annealed to generate a 28 bp double-stranded insert and cloned into p1495. Sequence-defined second-guide plasmids were verified by whole-plasmid sequencing before use in downstream library assembly.

### Background suppression during module construction

FORGE module vectors were designed with negative-selection sites to reduce background from empty vectors, vector re-ligation, or incorrect assembly products. Following Golden Gate assembly, reaction products or recovered plasmid pools were subjected to I-SceI (NEB, R0694S)- and I-CeuI (NEB, R0699S)-mediated background suppression when applicable^30^. Because correctly assembled modules disrupt the negative-selection cassette, digestion selectively depleted undesired assembly products and enriched for correctly assembled guide, barcode, or second-guide modules.

### Transformation, titering, and bacterial expansion of module libraries

Golden Gate assembly products were transformed into high-efficiency competent E. coli. For each transformation, 2 µL of the Golden Gate reaction was transformed and recovered in 300 µL SOC medium. Transformation efficiency and library complexity were estimated by plating 30 µL of recovered cells and 30 µL of a 1:100 dilution. Colony counts from titration plates were used to estimate the total number of transformants and to determine the number of transformations required for each library.

Transformation scale was adjusted according to library size to maintain representation during cloning. In general, one transformation was used for libraries containing fewer than approximately 1,000 guides, two transformations for libraries of approximately 1,650 guides, three transformations for libraries of approximately 3,000 guides, and five transformations for libraries of approximately 5,000 guides. For bacterial expansion, transformants were plated at approximately 20,000 colonies per 15 cm plate. One transformation was typically expanded on 4–5 plates, two transformations on 8–10 plates, three transformations on approximately 15 plates, and five transformations on approximately 25 plates. Colonies were harvested by scraping and pooled to generate bacterial library stocks. Aliquots of the pooled scraped cells were stored as glycerol stocks for subsequent plasmid preparation and reuse.

### Preparation of plasmid DNA from reusable guide-module libraries

During workflow development, plasmid DNA prepared directly from bacteria scraped from 15 cm agar plates was found to inhibit downstream enzymatic reactions, consistent with carryover of agar or agar-associated contaminants during plate harvest. To minimize this inhibition, plasmid DNA for reusable guide modules was prepared from liquid outgrowths seeded from the archived scraped-cell glycerol stocks rather than directly from scraped plate material.

For reusable guide Modules 1–7, 10, and 11, 30 µL of the corresponding glycerol stock generated from the pooled scraped bacterial library was inoculated into 50 mL LB medium containing kanamycin. Cultures were grown for 18 h at 30°C, and plasmid DNA was then isolated by midi-prep (Zymo Research, D4201-A). These midi-prepped plasmid DNA preparations were used as the input DNA for PCR–NGS library validation performed at the UCLA Technology Center for Genomics & Bioinformatics (TCGB). The same DNA preparations were archived as reusable guide-module plasmid stocks for downstream assembly into screening acceptor vectors.

### Assembly of reusable modules into FORGE-CRISPR screening acceptor vectors

Reusable guide, barcode, and second-guide modules were designed for transfer into pForge-CRISPR screening acceptor vectors by BsmBI-mediated Golden Gate assembly (NEB, E1602S). Guide-only libraries were generated by assembling pForge-CRISPR-Module-Single-Guide libraries directly into compatible acceptor vectors. Guide-plus-barcode libraries were generated by combining pForge-CRISPR-Module-Paired-First-Guide libraries with pForge-CRISPR-Module-Paired-Second-Barcode libraries during acceptor-vector assembly. Guide-plus-second-guide libraries were generated by combining pForge-CRISPR-Module-Paired-First-Guide libraries with pForge-CRISPR-Module-Paired-Second-Guide constructs.

Reusable guide, barcode, and second-guide modules are designed for transfer into pForge-CRISPR screening acceptor vectors by BsmBI-mediated Golden Gate assembly. Guide-only libraries are generated by assembling single-guide modules directly into compatible acceptors. Guide-plus-barcode and guide-plus-second-guide libraries are generated by combining paired-first-guide modules with barcode or second-guide modules during acceptor-vector assembly. Representative acceptor assemblies generated in this study included guide-only CRISPRi, guide-plus-barcode CRISPRi, guide-plus-barcode all-in-one CRISPR knockout, and paired-guide CRISPRi configurations; the paired-guide library carried a fixed KRAS second guide. Comprehensive cellular benchmarking of these final screening formats was not performed in this version.

### Plasmid archiving and reuse of modular libraries

Reusable module libraries were archived at multiple stages of construction. Bacterial stocks generated from pooled scraped colonies were stored as glycerol stocks, and midi-prepped plasmid DNA from liquid outgrowths was stored as reusable module DNA. Because guide, barcode, and second-guide modules were generated independently of the final screening vector, archived module libraries could be reused to generate additional screening formats without re-amplifying the original synthetic oligonucleotide pools. Material recovery and process performance were tracked across the workflow, including PCR yield, cloning efficiency, transformation efficiency, plasmid recovery, and final library yield.

### Quality control of cloned libraries

Cloning quality was assessed at multiple stages before sequencing. PCR products were examined by agarose gel electrophoresis to confirm the expected amplicon size and to evaluate primer removal after cleanup. Transformation efficiency was assessed by colony titration, and total transformant numbers were compared with library complexity to ensure adequate coverage. Plasmid DNA yield was measured after midi-prep. For reusable guide Modules 1–7, 10, and 11, the midi-prepped DNA prepared from liquid outgrowths of scraped-cell glycerol stocks was used for PCR–NGS validation.

### Preparation of sgRNA library amplicons for next-generation sequencing

Plasmid DNA libraries containing sgRNA pools were analyzed by next-generation sequencing to determine sgRNA representation and distribution within each library. A two-step nested PCR strategy was used to amplify the sgRNA-containing region from plasmid DNA. The first PCR incorporated a random 8-nucleotide unique molecular identifier (UMI) to facilitate downstream control of PCR duplicates^7,31,32^.

For the first PCR, 10 ng of plasmid DNA was amplified in a 50 µL reaction using Phusion High-Fidelity DNA Polymerase (Thermo Scientific, F530). Each reaction contained 1× Phusion HF buffer, 200 µM dNTPs, 500 nM forward primer, 500 nM reverse primer, 10 ng plasmid DNA template, and 0.5 U Phusion DNA Polymerase. The forward primer contained a random octamer sequence to serve as a UMI. PCR cycling was performed using an initial denaturation at 98°C for 30 s, followed by 10 cycles of 98°C for 10 s, 55°C for 30 s, and 72°C for 30 s, with a final extension at 72°C for 10 min. Following the first PCR, residual primers were removed using AMPure XP beads (Beckman Coulter) according to the manufacturer’s instructions.

The second PCR was performed in a 50 µL reaction using Phusion High-Fidelity DNA Polymerase to add sample indexes and sequencing adapters. Each reaction contained 1× Phusion HF buffer, 200 µM dNTPs, 500 nM i5-indexed forward primer, 500 nM i7-indexed reverse primer, 0.5 µL of purified first-PCR eluate, and 0.5 U Phusion DNA Polymerase. The second PCR was performed for 30 cycles to generate the final sequencing amplicons. After amplification, PCR products were purified using AMPure XP beads according to the manufacturer’s instructions.

Purified amplicons from different sgRNA libraries were pooled according to the number of guides in each library to achieve approximately equal representation of each sgRNA in the final sequencing pool. The pooled libraries were quantified by Qubit fluorometry and assessed for size distribution using a TapeStation system. Libraries were sequenced using 2 × 150 bp paired-end sequencing on a NovaSeq platform (Illumina) at the UCLA Technology Center for Genomics & Bioinformatics (TCGB). Sequencing data were analyzed using custom R scripts to quantify sgRNA abundance and evaluate library representation.

### Primer sequences

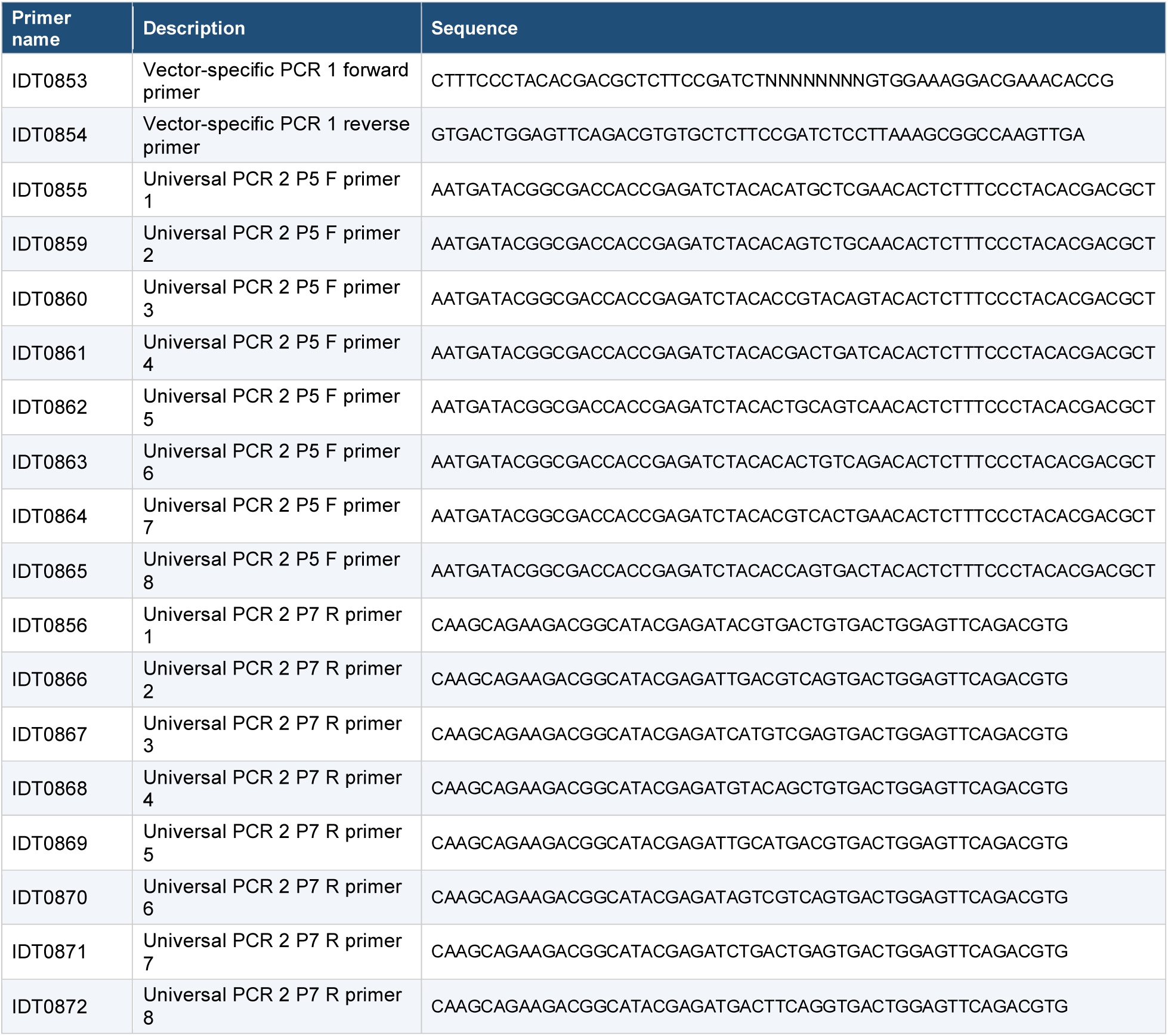

### Data and reagent availability

All FORGE-CRISPR modules and screening acceptor vectors have been deposited at Addgene under the deposit names listed in Table 1 (pForge-CRISPR-Module-Single-Guide, pForge-CRISPR-Module-Paired-First-Guide, pForge-CRISPR-Module-Paired-Second-Guide, pForge-CRISPR-Module-Paired-Second-Barcode, pForge-CRISPR-Acceptor-Cas9-GFP, pForge-CRISPR-Acceptor-PuroR-GFP, and pForge-CRISPR-Acceptor-PuroR-mScarlet). These constructs have been assigned Addgene catalog IDs #258796–#258802, listed per construct in Table 1. Annotated plasmid maps accompany each deposit, and the focused module guide lists summarized in Table 2 will be deposited alongside the plasmids in a future release. Processed data supporting this study will be deposited in a public data repository in a future release: per-guide read-count tables and per-module quality-control summaries (Figure 4); the cloning and material-economy records (Figure 5); guide lists for every constructed module; gene lists for every designed druggable-oncology module; annotated plasmid maps as GenBank files; and a README defining module and plasmid identifiers, guide-source versions, and dependency-classification rules.

## Author Contributions

J.-A.L., D.C., and M.P. contributed equally to this work. Conceptualization, J.-A.L., D.C., and M.P.; methodology, J.-A.L., D.C., and M.P.; software, D.C.; investigation, J.-A.L., D.C., and M.P.; formal analysis, J.-A.L., D.C., and M.P.; writing – original draft, J.-A.L., D.C., and M.P.; writing – review and editing, all authors; supervision, S.M.D. and J.M.L.; resources, S.M.D. and J.M.L.; funding acquisition, S.M.D. and J.M.L. All authors read and approved the final manuscript.

## Acknowledgments

Flow cytometry was performed in the UCLA Jonsson Comprehensive Cancer Center (JCCC) and Center for AIDS Research Flow Cytometry Core Facility, supported by National Institutes of Health awards P30 CA016042 and 5P30 AI028697, and by the JCCC, the UCLA AIDS Institute, the David Geffen School of Medicine at UCLA, the UCLA Chancellor’s Office, and the UCLA Vice Chancellor’s Office of Research.

